# Time-restricted feeding prolongs lifespan in *Drosophila* in a peripheral clock-dependent manner

**DOI:** 10.1101/2020.09.14.296368

**Authors:** Daniel Cabrera, Michael W. Young, Sofia Axelrod

## Abstract

Time-restricted feeding/eating (TRF/TRE) – limiting not the amount of food but the daily time window of food intake – is a dietary intervention that has been shown to improve health markers in model organisms and humans, but whether these benefits translate into positive effects on aging and longevity is not clear. We demonstrate here that TRF robustly prolongs lifespan in the short-lived genetically tractable model organism *Drosophila melanogaster*. Median TRF lifespan extensions range between ∼10% and ∼50% dependent on sex, reproductive status, TRF duration, and genotype. TRF’s positive effect on longevity is independent of food intake and at least in part relies on a functioning circadian clock: TRF benefits on longevity are abolished in arrhythmic *per*^*0*^ and *tim*^*01*^ mutants as well as in constant light, suggesting that timed feeding acts as a zeitgeber partitioning eating and associated metabolic processes into certain phases of day and night. TRF-mediated longevity extension is unaffected in flies whose neural circadian clocks have been abolished genetically, pointing towards peripheral clocks as the target of TRF mediating lifespan extension.

## Introduction

Aging, the deterioration of organismal function with time (Flatt and Schmidt, 2009), has been observed in almost all tested life forms (Ackermann et al., 2003; Lewis and Buffenstein, 2016), and is evident at the molecular, cellular, tissue and organismal levels. Researchers identified multiple correlates of aging including genomic instability, telomere attrition, epigenetic alterations, loss of proteostasis, deregulated nutrient sensing, mitochondrial dysfunction, cellular senescence, stem-cell exhaustion and altered intercellular communication (López-Otín et al., 2013). The causes for these changes are not clear, but candidate processes include macromolecular damage induced by reactive oxygen species (ROS), time-dependent and stochastic changes in levels of longevity-regulating molecules, circadian clock deterioration and hypertrophy (Gems and Partridge, 2013; Kenyon, 2010; Mattis and Sehgal, 2016; Musiek and Holtzman, 2016). The relative importance of both causes and hallmarks of aging and their relationships with each other remain unclear.

Aging is controlled both by intrinsic as well as environmental factors, including diet. First discovered in rats during the Great Depression, scientists found that reducing food availability, surprisingly, did not shorten but extended lifespan (McCay et al., 1989). Dietary restriction (DR) was later found to prolong longevity in many species from yeast to primates and improves health markers in humans, suggesting an evolutionarily conserved process (Chaix et al., 2019b; Colman et al., 2009; Partridge et al., 2005; Walker et al., 2005). Forward genetic screens probing the genetics of aging revealed that, indeed, two interconnected nutrient-sensing pathways, the insulin/insulin-like growth factor 1 (IGF-1) signaling (IIS) pathway and the target of rapamycin (TOR) pathways are required for DR’s effects on longevity (Kenyon, 2010). DR, at least in part, acts on the IIS and TOR pathways to change cellular transcriptional programs, involving the S6 kinase and FOXO transcription factors. Downstream events include upregulation of a transcriptional program making cells more resistant to molecular damage and curtailing growth. In addition to DR including caloric restriction and restricting uptake of certain nutrients or amino acids, other dietary interventions have been shown to have positive effects on aging: intermittent fasting, which includes various protocols of feeding or eating and fasting for longer than 24h (Patterson and Sears, 2017), and time-restricted feeding (TRF), where food intake is restricted daily to certain hours of the day, but total dietary intake is not (Chaix et al., 2019b). In *Drosophila*, TRF has been shown to attenuate age-related cardiac decline during aging (Gill et al., 2015), as well as increase sleep and muscle function (Villanueva et al., 2019). In mice, TRF lowers blood cholesterol and sugar levels, reduces body weight, inflammation and dysbiosis, while increasing energy expenditure, motor control and endurance (Chaix et al., 2019b). Human studies confirmed TRE’s potential as a health-promoting intervention and showed that TRF can help improve a number of metabolic markers including weight, body fat, blood pressure, energy intake and endurance, in particular in individuals at risk for metabolic syndrome (Chaix et al., 2019b; Wilkinson et al., 2020). TRF is distinct from DR as its health benefits do not seem to require a reduction in caloric intake (Gill et al., 2015; Hatori et al., 2012; Mitchell et al., 2019). TRF’s mechanism of action is not well understood, however it seems to increase the amplitude of circadian rhythms of various physiological parameters, which normally flatten during aging, presenting a possible mechanism of action (Manoogian and Panda, 2017).

While TRF’s effects on health and aging have been described, it is currently unknown whether TRF also extends longevity (Francesco et al., 2018), although single meal-fed as well as calorically restricted mice exhibit lifespan extensions that are correlated with self-imposed daily fasting (Acosta-Rodríguez et al., 2017; Mitchell et al., 2019). As TRF effects, like other dietary interventions, seem to be conserved across evolution, it is useful to employ short-lived genetically tractable model organisms to address this question. In the present study we used *Drosophila melanogaster*, which live only three months, therefore uniquely enabling us to do a high dimensional multi-parameter analysis to investigate whether TRF has an effect on longevity, and what the relevant biological factors and underlying mechanisms are.

## Results

### TRF prolongs lifespan in *Drosophila*

To assess whether TRF affects longevity, we exposed females, males and virgin *Drosophila* from an isogenic wild-type strain to a diurnal 12 hour food, 12 hour agar regimen in regular light:dark (LD) cycles for their entire lifespan (Fig. 1A), allowing the flies to eat during the day and have access to water-containing agar during the night to avoid desiccation. As population density, genetic background and food composition all independently affect longevity (Ivanov et al., 2015; Joshi and Mueller, 1997; Skorupa et al., 2008), each of these parameters was kept constant across all experiments unless explicitly noted (see Materials and Methods for more details). TRF significantly extended median lifespan in mated female flies (called ‘flies’ from here on unless stated otherwise) from 49 days to 60 days on average (+18%), but not in mated males or virgin females (Fig. 1B,C, Table 1 and Supp. Fig. 1). Short-term TRF/TRE has been shown to be beneficial for cardiac health and sleep in *Drosophila* (Gill et al., 2015) and metabolic markers in rats and humans (de Goede et al., 2019; Hutchison et al., 2019; Jamshed et al., 2019; Ravussin et al., 2019). In order to determine whether shorter-than-lifelong TRF still positively affects longevity, we tested 10, 20, 30, 40 and 50 days of TRF compared to ALF. TRF showed a dose-response effect depending on TRF duration, with a minimum duration of 20 days to achieve longevity extension (Fig. 1D, left panel, and table 1). To assess whether TRF in the second half of life is still beneficial, we exposed flies to TRF from days 30-60 of age. TRF in the second half of the flies’ life significantly extended longevity (Fig. 1D, right panel), showing that TRF is beneficial independently of the age at which it is started.

**Table 1.**
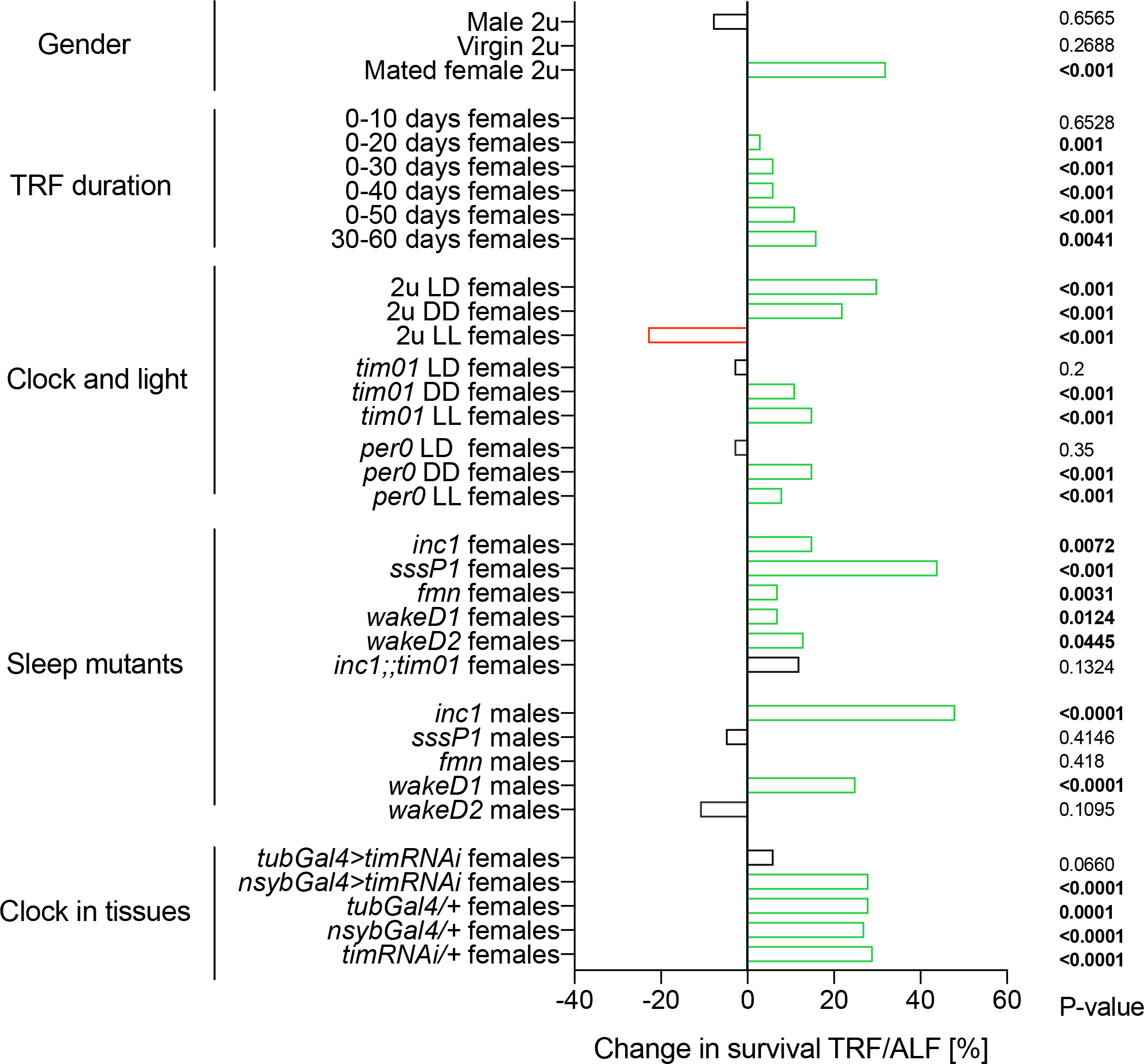
Summary of described TRF effects. Left column denotes experimental groups elucidating the role of different parameters on TRF effects. Bar chart indicates positive (green), neutral (black) or negative (red) effects of TRF on longevity. P-values in bold on the right indicate significant changes.

**Fig 1.**
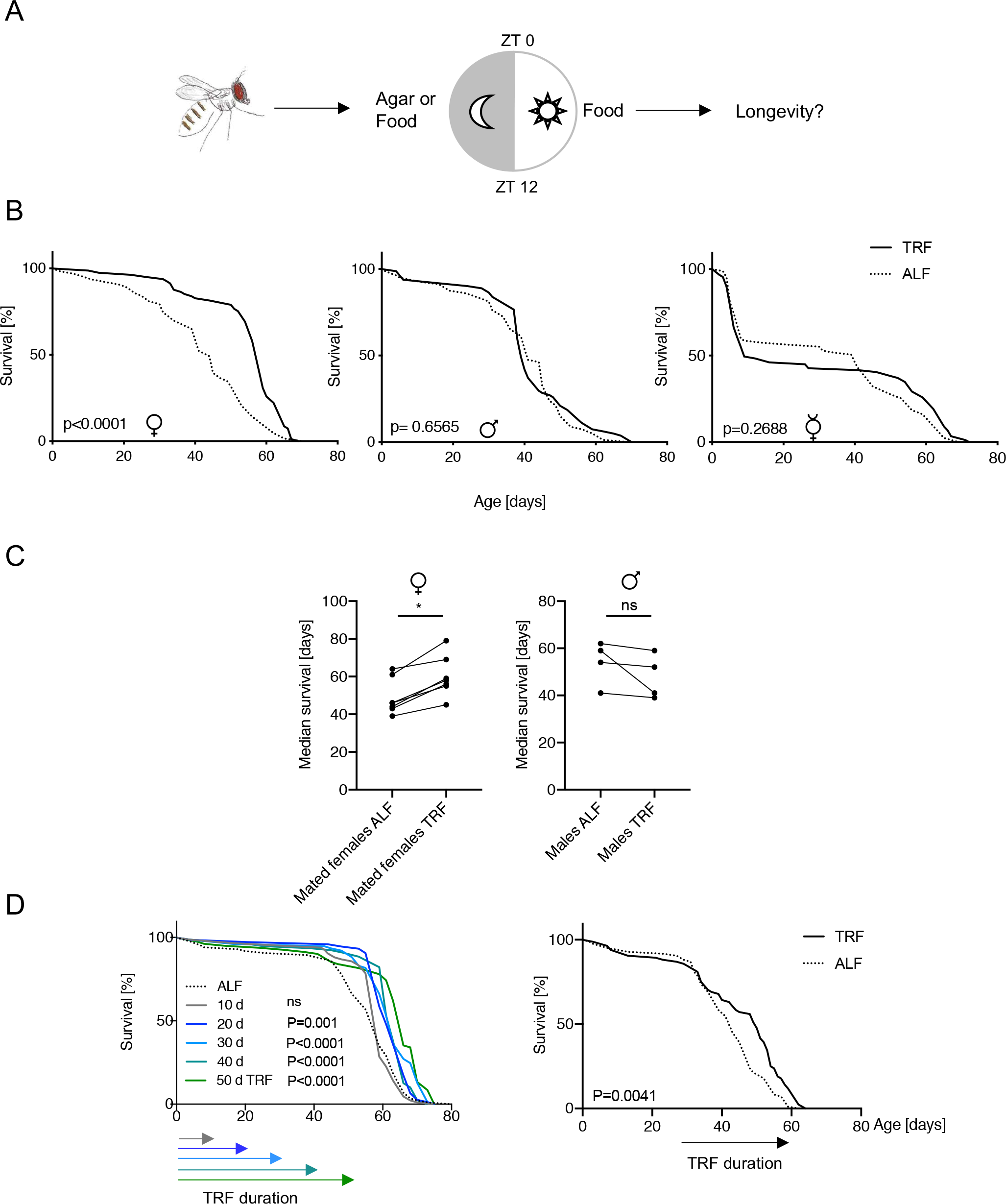
Time-restricted feeding (TRF) prolongs longevity in wild-type *Drosophila* females. (A) A simplified diagram of the TRF method used. Flies were transferred at zeitgeber time (ZT) 12 from food to agar and ZT 0 from agar to food to restrict food intake to 12 h during the day (TRF group) or from food to food as a control (ALF). Fly survival was assessed daily. (B) TRF significantly extends lifespan in mated females, but not in mated males or virgins (ALF n=63,80,87, TRF n=81,83,89, respectively). (C) TRF reproducibly extends longevity in (left) mated females (ALF ns=63, 59, 79, 95, 116, 82, 153. TRF ns=81, 69, 67, 75, 105, 84, 142) but not (right) males (ALF ns=80, 34, 77, 95, TRF ns=83, 38, 83, 90) across different experiments (See Supp. Fig. 1 for additional survival plots). *=P≤0.05 (D) TRF acts in a duration-dependent manner. (Left) Mated females underwent TRF for 10, 20, 30, 40, 50 days (ALF n=86, 10, 20, 30, 40, 50d, TRF ns=84, 75, 77, 79, 82, respectively) or (Right) from 30–60 days of age (ALF n=84, TRF n=84). (Left) Lifespan extension with TRF is observed with a minimum duration of 20 days and increases with longer TRF. (Right) TRF in the second half of the flies’ life significantly prolongs lifespan.

### Partial restoration of normal lifespan through TRF in short-lived sleep mutants

*Drosophila* longevity is affected by a number of factors including temperature, light conditions, genetic background, and sleep (Bushey et al., 2010; Lamb, 1968; Malick and Kidwell, 1966; Sheeba et al., 2000). Most *Drosophila* sleep mutants exhibit shortened lifespans (Koh et al., 2008; Kume et al., 2005; Liu et al., 2014; Rogulja and Young, 2012; Stavropoulos and Young, 2011). To test whether TRF effects are restricted to normal-lived wild-type flies or also promote longevity in mutations affecting sleep and, in most cases, lifespan, we exposed the short-lived sleep mutants *sss*^*P1*^, *inc*^*1*^, *wakeD1, wakeD2*, as well as *fumin*, which had been described as having normal lifespan, to TRF or ALF for life. TRF significantly extended lifespan in all tested mated female sleep mutants (Fig. 2A and Table 1), with a lifespan extension of 50% for *sss*^*P1*^, 14% for *wakeD2*, 13% for *inc*^*1*^, 12% for *wakeD1* and 9% for *fumin*. Interestingly longevity in some male sleep mutants was also aided by TRF (Fig. 2C, Table 1), in contrast to wild-type flies, where TRF did not extend males’ lifespan (Fig. 1B, Table 1, Supp. Fig. 1B). These data illustrate that TRF effects are not limited to normal-lived wild-type flies but promote longevity to varying degrees in other genotypes, specifically short-lived sleep mutants. Because TRF works on some male strains, we conclude that it is, in principle, possible to prolong longevity in both females and males by using TRF. Increased sleep duration does not appear to be required for TRF’s effect on longevity: While we confirmed the previously reported TRF-induced increase in total daily sleep in males (Gill et al., 2015), data not shown), these flies do not exhibit a lifespan increase (Fig. 1B, Supp. Fig. 1B). In contrast, TRF-treated females flies do not show a change in total daily sleep, although their daytime sleep is increased compared to ALF flies (Supp. Fig 2). Importantly, TRF effects on sleep are absent in the sleep mutant *inc*^*1*^ (Supp. Fig 2), although longevity effects are not (Fig. 2), and conversely *per*^*0*^ mutants show increases in nighttime sleep (Supp. Fig. 2) but no longevity effects (Fig. 4), suggesting that TRF improves longevity independent of effects on sleep.

**Fig 2.**
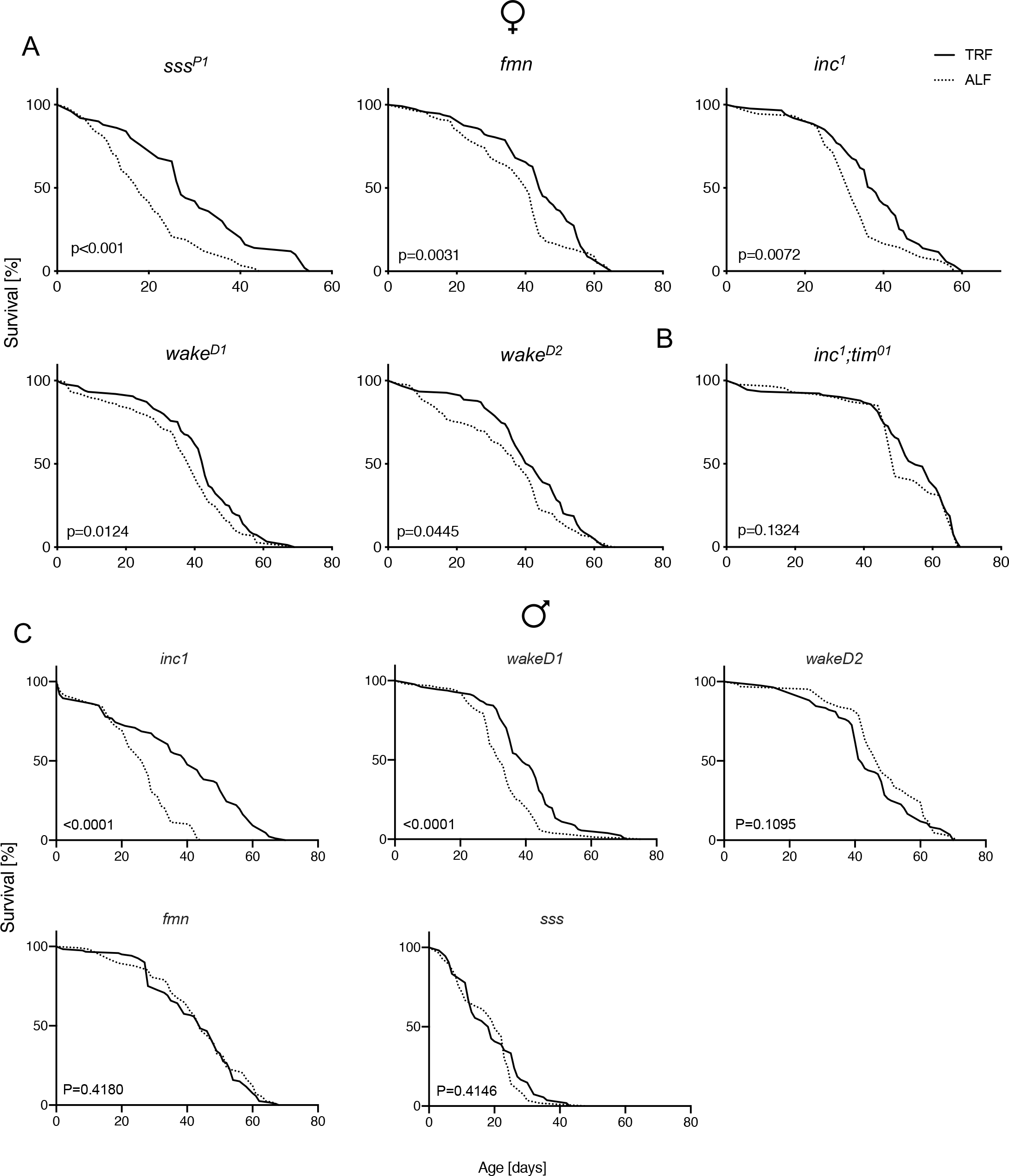
TRF improves longevity in sleep mutants with shortened lifespans in a clock, sex and genotype-dependent manner. Female or male flies were kept on TRF or ALF for the duration of their lives. (A) All indicated genotypes show TRF-mediated lifespan extension. *Inc*^*1*^ (ALF n=91, TRF n=87), *sss*^*P1*^ (ALF n=58, TRF n=50), *fmn* (ALF n=136, TRF n=113), *wakeD1* (ALF n=144, TRF n=149), and *wakeD2* (ALF n=151, TRF n=123). (B) *tim*^*01*^ null mutation reverses TRF-mediated lifespan extension in *inc*^*1*^ flies (ALF n=87, TRF n=91). (C) TRF promotes lifespan extension in *inc*^*1*^ (ALF n=88, TRF n=86) and *wakeD1* (ALF n=155, TRF n=127), but not *wakeD2* (ALF n=63, TRF n=84), *fmn* (ALF n=114, TRF n=120) and *sss*^*P1*^ (ALF n=57, TRF n=54) flies.

**Fig 3.**
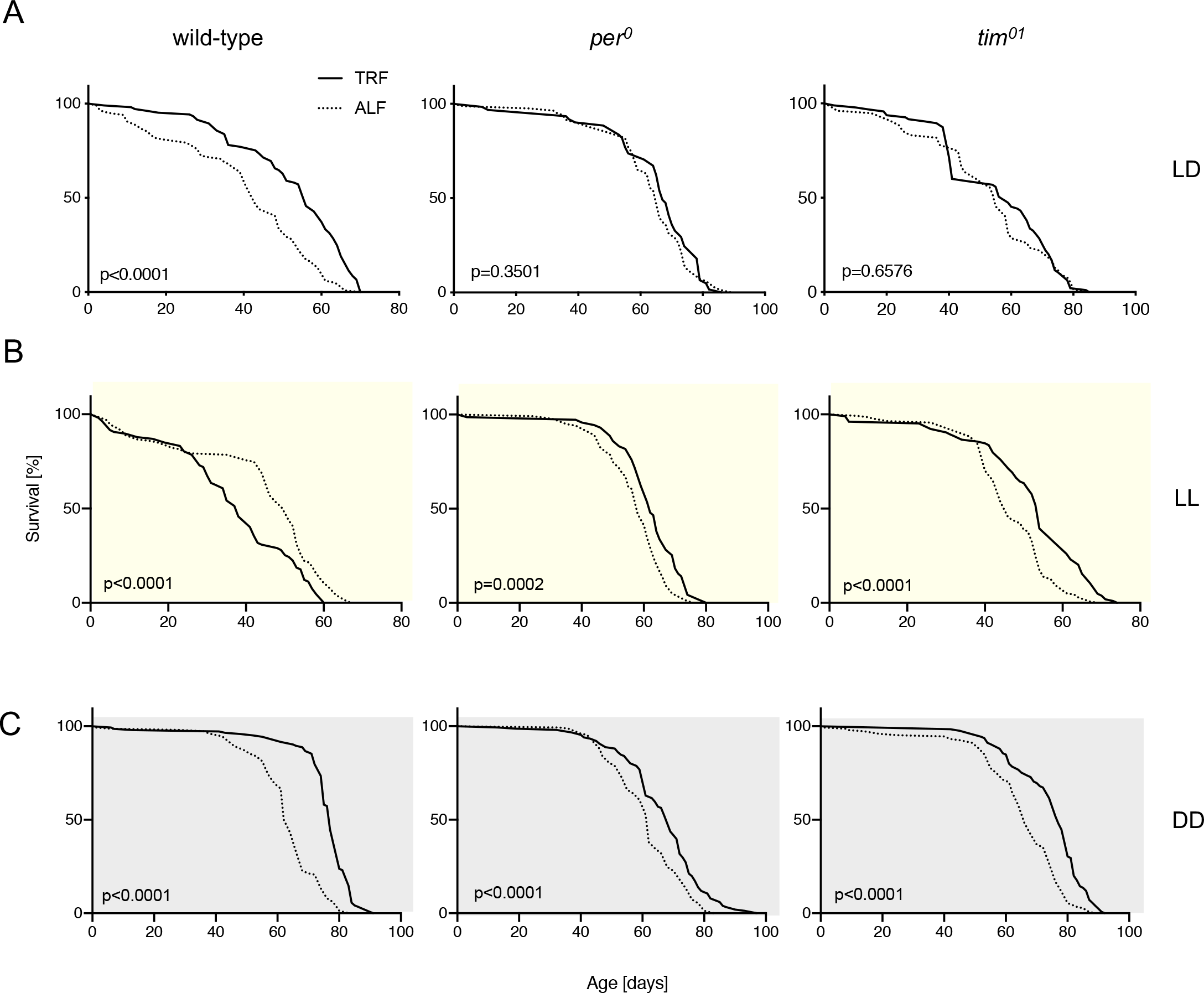
Clock mutations reverse TRF-mediated lifespan extension in a light-dependent manner. Female wild-type, *per*^*0*^ or *tim*^*01*^ flies were kept on 12:12 TRF or ALF for the duration of their lives. (A) In 12:12 light:dark (LD) wild-type (left), per (middle) and tim mutations (right) (ALF n=116,83,78, TRF n=105,60,95, respectively) abrogate TRF-mediated lifespan extension (left). (B) In constant light, TRF no longer extends longevity in wild-type flies, but prolongs longevity in *per*^*0*^ and *tim*^*01*^ (ALF n=135,119,137, TRF n=107,71,104, respectively) mutants. (C) In constant darkness, TRF promotes longevity extension in wild-type, *per*^*0*^ and *tim*^*01*^ (ALF n=150,147,150, TRF n=143,125,151, respectively).

**Fig 4.**
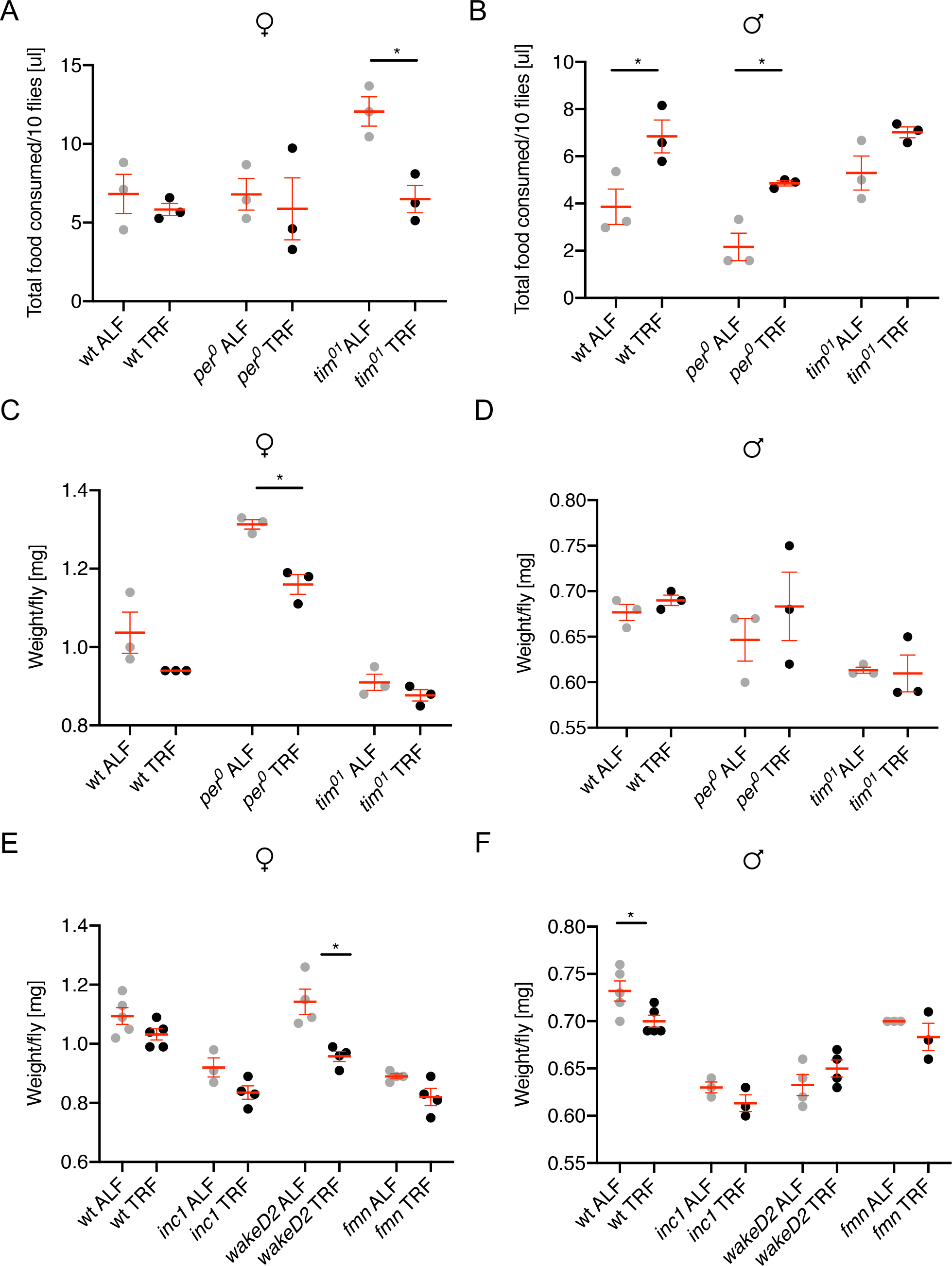
Impact of TRF on feeding and bodyweight. (A) Female or (B) male wild-type, *per*^*0*^ and *tim*^*01*^ flies underwent 14 days of TRF or ALF in 12:12 LD. Food intake was measured in groups of 10 flies for 12 hours (TRF group) or 24 hours (ALF group) using the capillary feeder assay (CAFE). Each point represents food consumption of 10 flies. (C) Female or (D) male wild-type, *per*^*0*^ and *tim*^*01*^ flies underwent 14 days of TRF or ALF in 12:12 LD and were weighed in groups of 10 at ZT 10. (E) Female or (F) male *inc*^*1*^, *wakeD2* and *fmn* underwent 30 days of TRF or ALF in 12:12 LD and were weighed in groups of 10 at ZT 10. Each point represents the weight of 10 flies divided by 10. Mean and SEM are indicated in red. Mann-Whitney testing assessed statistical significance. *=P≤0.05

### TRF effects are clock and light dependent

The circadian clock consists of molecular machinery present in most cells and conserved across evolution allowing organisms to align physiological parameters with daily environmental cycles by providing temporal organization to biological processes and behaviors (Patke et al., 2019). TRF effects on cardiac health have been shown to be dependent on the circadian clock (Gill et al., 2015). To investigate whether an intact molecular circadian clock is also necessary for TRF-mediated lifespan extension, we tested the arrhythmic *per*^*0*^ and *tim*^*01*^ clock mutants (Konopka and Benzer, 1971; Sehgal et al., 1994). TRF effects on longevity are absent in *per*^*0*^ and *tim*^*01*^ mutants (Fig. 3A), as well as in an *inc*^*1*^, *tim*^*01*^ double mutant (Fig. 2B), suggesting that a functioning circadian clock is required for TRF-mediated longevity effects. This conclusion is strengthened by TRF experiments conducted in constant light (‘LL’), which renders wild-type flies arrhythmic due to constant resetting of the molecular clock via timeless degradation (Myers et al., 1996; Zeng et al., 1996). In LL, TRF has the opposite effect compared to LD: it is detrimental to the flies’ longevity (Fig. 3B, left). In constant darkness, TRF benefits to wild-type flies persist (Fig. 3C), and TRF was also beneficial to the *per*^*0*^ and *tim*^*01*^ clock mutants, which saw longevity extension in constant light as well (Fig. 3B). These data suggest that TRF effects depend on a functioning circadian clock as well as on the light/dark cycle.

### TRF longevity benefits are not dependent on decreased food intake

Caloric (or dietary) restriction (DR) is a known mechanism to prolong lifespan in various model organisms (Colman et al., 2009; Partridge et al., 2005). To test whether TRF simply leads to DR, we measured the flies’ food consumption after 14 or 32 days of TRF or ALF. Utilizing the capillary feeder assay (CAFE (Ja et al., 2007)), we determined that TRF does not cause flies to eat less as TRF-treated female wild-type flies which exhibit lifespan extension under TRF consume similar amounts of food during the 12 hours they have access to food than ad libitum controls in 24 hours (Fig. 4A, Supp. fig. 3A), demonstrating that TRF is distinct from DR. In contrast to that, *tim*^*01*^ flies do exhibit a reduction in food intake under TRF (Fig. 4A), however that does not translate into longevity extension in our assay (Fig. 3A). To test how TRF affects the flies’ body weight, which is associated with different metabolic markers as well as aging in *Drosophila* and humans (Fontana and Hu, 2014; Gáliková and Klepsatel, 2018), we weighed flies in 3 or 4 groups of 10 after 14, 30 or 45 days of daily TRF or ALF. Although wild-type female flies show weight loss after 45 days of TRF (Sup. Fig 3B), 14 or 30 days of TRF don’t have a significant effect on their weight (Fig. 4C,E). Weight loss was also observed for some other genotypes (*per*^*0*^, *wakeD2*, Fig. 3 C,E, Supp. Fig. 3B) and for wild-type males after 30 days of TRF (Fig. 3F), however these conditions are not associated with a TRF-mediated longevity increase. Therefore, weight loss occurred independently of TRF-induced longevity effects suggesting that weight loss is not the primary mechanism leading to prolonged lifespan.

### TRF-mediated lifespan extension does not require a neuronal circadian clock

While most cells have molecular circadian clocks, peripheral tissues including muscles, liver and kidneys can receive time-of day information either directly from environmental cues or from a specific brain structure called the central or master clock (Patke et al., 2019). *Drosophila* locomotor rhythms and the sleep/wake cycle are controlled by so-called clock neurons in the central brain, a subset of which, along with the circadian neuropeptide pigment dispersing factor, have also been implicated in aging (Donlea et al., 2014; Umezaki et al., 2012). To elucidate, whether a functional neuronal clock is required for TRF effects on longevity, we used the pan-neuronal Gal4 driver *nsyb-Gal4* to knock down *tim* RNA using *UAS-tim-RNAi*, which renders flies behaviorally arrhythmic (Supp. Fig. 4). While ubiquitous *tim-RNAi* in all tissues using *tubulin-Gal4* phenocopies the abrogation of TRF effects observed in the *tim*^*01*^ null mutants (Figs. 4A, right panel and 5A), neuronal-specific RNAi left TRF effects intact (Fig. 5B), suggesting that nervous system-based circadian clocks are dispensable for TRF effects on longevity and that peripheral clocks can support TRF-mediated lifespan extension.

**Fig 5.**
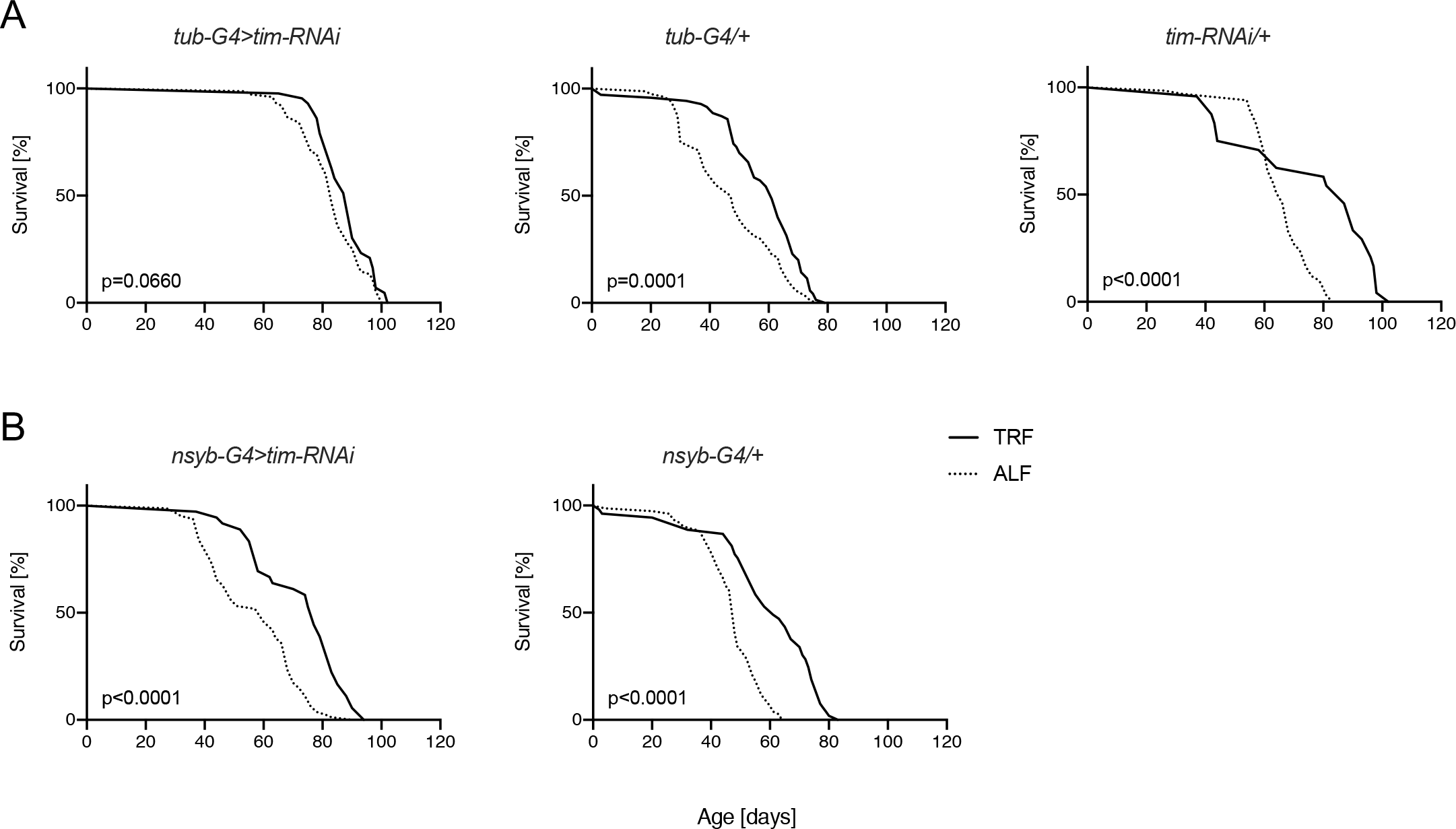
TRF extends longevity independently of neuronal clocks. (A) Ubiquitous knockdown of *tim* using *tim-RNAi* abrogates TRF-mediated longevity extension (left: knockdown (ALF n= 75, TRF n=43), middle: *tub-Gal4* driver control (ALF n=80, TRF n=70), right: *tim*-*UAS-RNAi* control (ALF n=75,80,67, TRF n=24)). (B) Left: Pan-neuronal *tim-RNAi* (ALF n=81, TRF n=36) has no effect on TRF-mediated longevity extension. Middle: *nsyb-Gal4* neuronal driver control (ALF n=77, TRF n=53)

## Discussion

TRF/TRE has garnered attention in recent years as a dietary intervention producing health benefits in humans and animal models, but TRF’s effect on longevity, as well as underlying principles of TRF benefits, have remained largely unknown. In the present study we demonstrate clock and light-dependent, yet neuron-independent, life extension by TRF.

### TRF as a novel life extension mechanism

Dietary Restriction (DR) – reducing the intake of calories or specific macronutrients – has been described as a powerful method to prolong longevity in yeast, *C. elegans, Drosophila*, mice and primates (Colman et al., 2009; Partridge et al., 2005). We observe that TRF-mediated longevity benefits are not dependent on reduced food intake, which is in line with previous studies in flies and mice showing that other TRF induced health benefits, too, are independent of DR (Gill et al., 2015; Hatori et al., 2012; Mitchell et al., 2019). While weight loss occurs in some conditions where TRF prolongs lifespan, it also occurs when lifespan is not prolonged and vice versa, suggesting that weight loss is not the primary cause for TRF’s longevity extension, but rather downstream of its (yet unknown) mechanism. In contrast to DR, TRF shows duration-dependent benefits starting at two weeks of TRF, (Fig. 1D), while DR effects on mortality rates have been shown to take hold after only 48h (Mair et al., 2003). In mice, DR seems to work only in young animals (Hahn et al., 2019), while in the present study TRF is equally effective in young and old animals. These data add additional evidence that TRF is mechanistically distinct from DR. On the other hand, DR, in some experimental paradigms, is hard to distinguish from TRF. Two recent studies attempted to compare TRF and DR in mice, however mice under DR self-imposed TRF and ate all food pellets in a narrow one to five hour window, effectively making DR indistinguishable from TRF (Acosta-Rodríguez et al., 2017; Mitchell et al., 2019). A novel timed food distribution assay will allow mouse researchers to distinguish these two dietary interventions (J. Takahashi, unpublished). Another dietary intervention promoting health benefits in various model organisms including *C. elegans, Drosophila*, mice and in humans is intermittent fasting (IF), where organisms are exposed to prolonged (>=24h) periods of fasting. While IF has been shown to improve metabolic markers (Patterson and Sears, 2017) and longevity by acting independently from the TOR pathway (Catterson et al., 2018) – a major target of DR –, it is unclear whether TRF acts through the same molecular pathways as IF. In the future, it will be interesting to better tease apart mechanistic differences and commonalities between different dietary interventions on aging.

### Sex differences in TRF effects

We observe that TRF effects on longevity are more universal across genotypes in females than in males. TRF extended median lifespan in all female genotypes tested, while in males three different mutant strains but not wild-type flies showed significant improvements with our assay (Fig. 2 and Table 1). While historically studies across most areas of biological research, including aging, have used only males, sex differences are emerging as an important research area in its own right. Male and female humans and model organisms including *Drosophila* age differently, with females generally living longer, however this relationship is neither universal nor fully understood, and highly dependent on factors including genetics, reproductive status and the environment, including diet (Austad and Fischer, 2016). Reproductive status, in addition to overall lifespan, also seems to affect survival dynamics on a population level. Virgins seem to follow a different survival trajectory than males and mated females (Fig. 1 B), with the absence of a delay of deaths occurring in the flies’ early stage of life. This deviation from the classic “S-shaped” survival plot seems to be a feature of virgin *Drosophila*, as previous studies also reported similar survival dynamics (Markow, 2011; Villanueva et al., 2019). In flies, TRF has been shown to have beneficial effects on sleep in males ((Gill et al., 2015), female data not shown in that study), and improve cardiac health and muscle function in both sexes (Gill et al., 2015; Villanueva et al., 2019). Previous work examining the effect of TRF on longevity in both sexes found no benefit, in contrast to the data presented here. The difference is likely to come from the reproductive status of flies used: while the former study appears to have used virgin males and females, we found that in wild-type, only mated females respond with lifespan extension upon TRF treatment, a condition not tested in the former study (Villanueva et al., 2019). While we do not know why only mated females show a longevity increase with TRF, various considerations could play a role in mated females’ differential response. Female flies are larger than males, exhibit differences in their metabolism and are less sensitive to starvation (Austad and Fischer, 2016) and data not shown). Females undergo a metabolic shift upon mating to maximize egg laying, which alters their food composition preference and aging trajectory (Lee et al., 2008). Finally, females exhibit important differences to males in their circadian clock and locomotor behavior, in particular after mating (Fujii et al., 2007; Helfrich-Förster, 2000; Isaac et al., 2010). It will be interesting to find out which aspect(s) of female physiology makes them more amenable to TRF. Our finding that TRF does, in principle, work in males, too, suggests that adjusting TRF duration, phase or food composition might create conditions for a more universal male responsiveness.

### TRF, the circadian clock, and light regimens

We observe that TRF-mediated longevity extension is, under certain conditions – like DR (Katewa et al., 2015; Ulgherait et al., 2016) – dependent on an intact circadian clock, while under certain lighting conditions, TRF effects can be clock-independent. In LD cycles, the clock mutants *per*^*0*^ and *tim*^*01*^ do not show extended longevity in response to TRF when compared to ALF, and constant light, which causes arrhythmicity in wild-type animals, also abolishes lifespan extension by TRF, suggesting circadian clock dependency. Interestingly, constant conditions - DD or LL - allow clock mutants to benefit from TRF, which points towards clock-independent TRF benefits as previously described (Chaix et al., 2019a; Xu et al., 2011), and light as a secondary parameter. Light:dark cycles present a stimulus organizing diurnal rhythms in clock mutants independently of a circadian clock, and induce a gene expression profile distinct from yet partially overlapping with that driven by circadian clocks (Wijnen et al., 2006). Timed feeding on the other hand, is a zeitgeber which can increase circadian rhythms in *Drosophila* metabolic tissues with or without an intact clock (Xu et al., 2011) and increase the amplitude of metabolic parameters and sleep/wake cycles in clock-less mice (Chaix et al., 2019a), or, if presented at an unnatural time, negatively affect other physiological parameters (Xu et al., 2011). Our observations could be explained by a model in which the circadian clock, light, and food impulses interact in a complex manner: LD-and LL-triggered gene expression programs differ between wild-type and clock mutants. Gene expression programs induced by LD in clock mutants and LL in wild-type flies oppose beneficial TRF effects, while LD in wild-type and DD in all genotypes aligns with TRF effects promoting longevity. Further studies are required to fully understand this interaction. Light itself has been shown to play an important role in aging. Both light regimen and color composition of light differentially affect aging in *Drosophila* (Nash et al., 2019; Sheeba et al., 2000), so it will be interesting to see how light-induced molecular pathways contribute to TRF effects. While the adaptive significance of the circadian clock has been demonstrated in various forms of life including fruit flies and humans (Patke et al., 2019), the role of clock genes in longevity is more complex. Surprisingly, although sleep duration mutants in *Drosophila* have strong effects on lifespan, circadian clock mutants in *Drosophila* and mice do not show large lifespan deficits (Dubrovsky et al., 2010; Klarsfeld and Rouyer, 1998; Kondratov et al., 2006), or under certain conditions even prolong lifespan (Ulgherait et al., 2020). Future studies will show how the life-prolonging and life-shortening aspects of food, light and the clock interact on a mechanistic level to determine an animal’s lifespan.

We observe that the clock gene *tim*^*01*^ is not required in neurons for TRF’s longevity effects, indicating that peripheral circadian clocks are sufficient for TRF effects. This finding is in line with previous results in mice showing that timed feeding acts on peripheral clocks (Manoogian and Panda, 2017). TRF increases the circadian amplitude of various metabolic factors, both in the mouse liver and in the fly fat body (the fly liver), while circadian rhythms in the brain remain unchanged (Xu et al., 2011). Furthermore, in mice, food out of phase with the sleep/wake cycle can create circadian dissonance within the body, with liver and brain clock operating at different phases (Damiola et al., 2000). Future research will aim at identifying the organ(s) where a functional clock is required, as well as pinpointing the molecular pathways mediating longevity benefits through tissue-specific clock function.

### Theories of aging

Two theories of aging postulate that aging is either the accumulation of molecular damage after reproduction has been accomplished - theory of the “disposable soma” (Kirkwood, 1977), or the result of hyperplastic processes - theory of the “bloated soma”, or hyperfunction theory, where hyperplasia and hypertrophy drive pathologies late in life (Gems and Partridge, 2013). TRF-mediated lifespan extension could affect aging via both categories: circadian clocks have been shown to increase detoxifying xenobiotic metabolism and resistance to molecular damage from reactive oxygen species (Gachon et al., 2006; Rakshit and Giebultowicz, 2013), suggesting that TRF might ameliorate aging by reducing the accumulation of molecular damage. On the other hand, our study shows that TRF flies lose weight, reducing biomass and potentially lessening hyperplasia and hypertrophy during aging, supporting the bloated soma theory. Future studies should reveal the exact molecular mechanism of TRF-mediated lifespan extension, potentially creating novel therapeutic targets for anti-aging and age-related pathologies.

## Methods

### *Drosophila* strains

Isogenic Iso1CJ (Yin et al. 1994) was used as wild-type. Circadian mutants *per*^*0*^ and *tim*^*01*^ (Konopka and Benzer, 1971; A. Sehgal et al., 1994) were used to study the effects of TRF on clock mutants. *UAS-timRNAi* and *tub-Gal4* are from the Bloomington stock center, #29583 and #5138, respectively. *nsyb-Gal4* is a lab stock. Sleep mutants *inc1* (Stavropoulos and Young, 2011), *fumin* (Kume et al., 2005), *sss*^*P1*^ (Koh et al., 2008), and *wakeD1* and *wakeD2* (Liu et al., 2014) were used to assay TRF on sleep mutants. All mutants were recently (<2 years) backcrossed to wild-type flies for at least 5 generations.

### Longevity Assay

Animals were collected after eclosing and allowed to mate and age for 2 days in vials containing fly food composed of 0.6% (w/v) agar (Mooranger Inc #41004), 5.25% (w/v) yellow cornmeal (VWR #IC90141180), 2.2% (w/v) yeast (Lesaffre Yeast Co. #73050), 5% (v/v) molasses (Woolco Foods Inc. #595650), 0.625% (v/v) tegosept (30% hydrobenzoic acid (Sigma #H5501) in 95% ethanol), 0.66875% (v/v) propionic acid (VWR #JTU330-9). Subsequently, animals were lightly anesthetized with CO_2_ and separated into vials with animals of the same sex, which were kept in LD cycles at 25°C. As population density affects longevity and crowding shortens lifespan, we placed 30 flies in each vial (Joshi and Mueller, 1997). Flies on TRF were manually transferred to vials with 1.1% agar at the end of the light part of the LD cycle, and transferred back to standard cornmeal at the beginning of the light part of the LD cycle. Animals on ALF were transferred daily to fresh food after initial experiments did not show a longevity difference between 12h and 24h ALF transfers (not shown). The number of dead animals was recorded every day. Experiments in constant light conditions were carried in an incubator at 25°C and lights on for the entirety of the experiment. Experiments in constant darkness were performed in a dark incubator and animals were handled with a red light in the dark room for scoring of survival and transfer to new vials. We used at least three technical replicates per condition in each experiment, totaling 90 flies per condition, which is in line with standards in the field, e.g. (Castillo-Quan et al., 2019). Masking was used during data collection and analysis. Survival Log-rank (mantel-cox) tests were performed in Graphpad Prism to compare survival of TRF v ALF groups.

### Sleep and Circadian Analysis

Single age matched animals from TRF experiments in LD were loaded into glass tubes containing standard cornmeal and assayed for four days in LD for sleep measurement and for five days in DD for rhythmicity using DAM5 monitors (Trikinetics). Locomotor data were collected in 1 min bins, and a 5 min period of inactivity (Huber et al., 2004; Shaw et al., 2000) was defined as sleep. Sleep was calculated using custom software in Python (Axelrod S. unpublished). Dead animals were excluded from analysis either by the software or by visual inspection. Mann-Whitney tests were performed to compare sleep between TRF and ALF groups using Graphpad Prism.

### Fly weight

To measure fly weight, we allowed all fly groups (ALF and TRF) to remain on fresh food for approximately 10 hours. Flies were separated into groups of 10 flies and weighted on a balance (Mettler AT20). Data reported are for triplicates or quadruplicates of the average weight of 10 flies. Mann-Whitney tests were performed to compare weight between TRF and ALF groups using Graphpad Prism.

### Feeding assay

We measured feeding in adult flies using the CApillary FEeder (CAFE) Assay (Diegelmann et al., 2017; Ja et al., 2007). 1-2 days old animals were picked and placed in LD at 25 C. A course of 14 or 32-day TRF was given to the TRF groups receiving food only from ZT 0-12 when lights are on, compared to ALF with constant food in equal conditions. After 14 or 32 days of TRF, 10 flies were placed in a vial covered with a cotton swab and a capillary tube with food and each measurement was done in triplicates. Animals were placed in plastic vials with 1.1% agar and liquid food containing 5% sucrose were delivered using borosilicate capillary tubes. The capillary was assessed every three hours for the length of food change and food was reloaded. For each experiment, a standard curve was generated by determining the change in capillary food length for different food volumes. As a control for evaporation, the measurements were reported as the difference between vials with flies and food to vials with food and no flies.

## Acknowledgements

The authors thank S. Lincoln for help with TRF flips and longevity assays, S. Jordan for helpful feedback on the manuscript and R. Jackson, A. Sehgal, M. Wu, and the Bloomington stock center for fly stocks.

## Author contributions

S.A. and M.W.Y conceived of the project. S.A., D.C. and M.W.Y designed experiments. D.C. performed experiments with guidance from S.A. and M.W.Y. and occasional help from S.A.. D.C. and S.A. analyzed the data, which were discussed by all authors. S.A. wrote the manuscript with input from all authors.

## Data and materials availability

The data reported are presented in the main paper and the supplementary materials.

---

**Supp. Fig 1.**
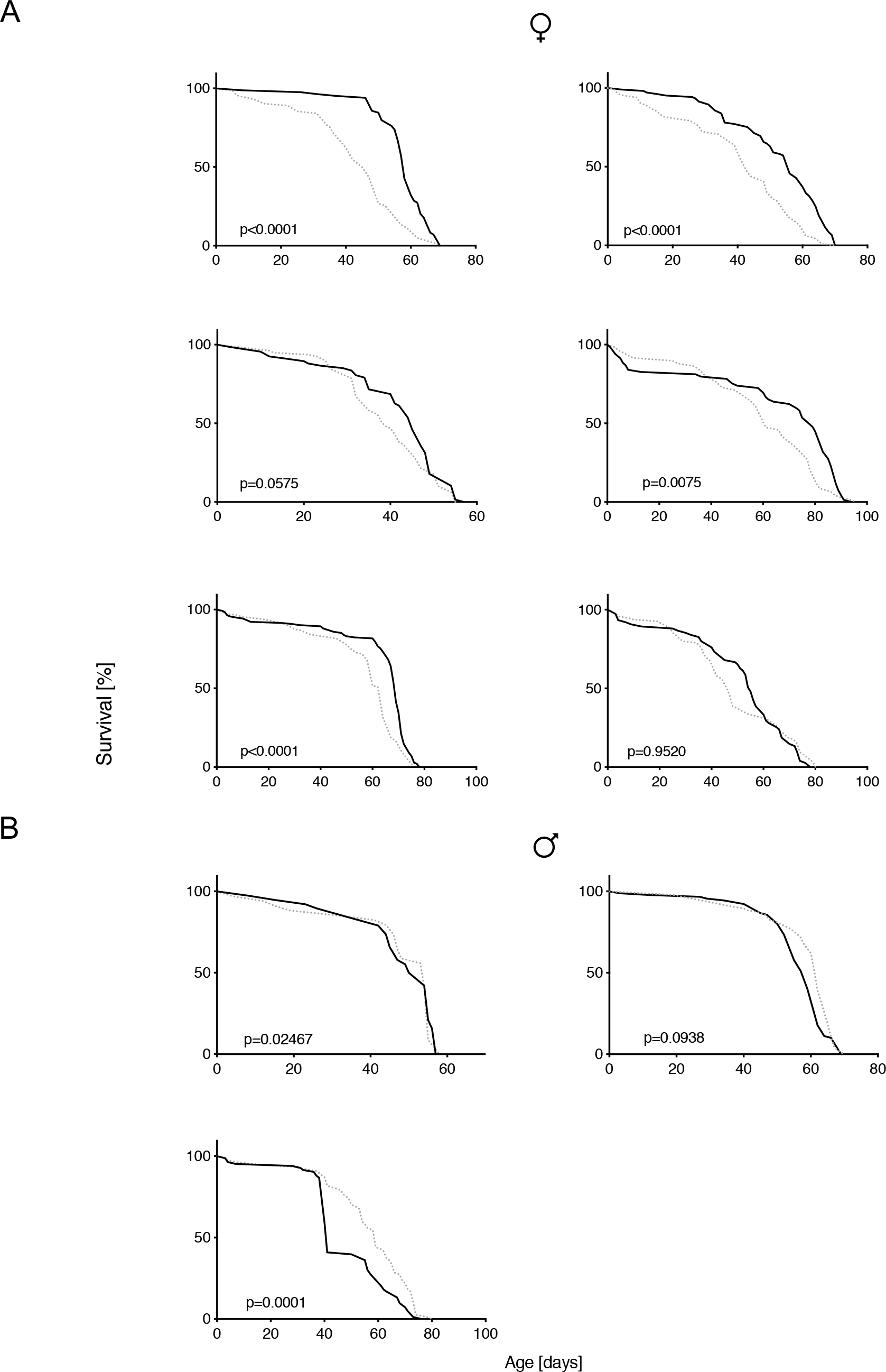
Time-restricted feeding (TRF) prolongs longevity in wild-type *Drosophila* females. Shown are individual experiments summarized in Fig. 1C. Flies were transferred at zeitgeber time (ZT) 12 from food to agar and ZT 0 from agar to food to restrict food intake to 12 h during the day (TRF group) or from food to food as a control (ALF). Fly survival was assessed daily. TRF significantly extends lifespan in mated females (A), but not in mated males (B). (A) Survival plots for TRF on females. n(ALF,TRF)=(82,84), (116,105), (79,67), (59,69), (153,142), (95,75), respectively. (B) Survival plots for TRF on males. n(ALF,TRF)= (34,38), (95,90), (77,83), respectively.

**Supp. Fig 2.**
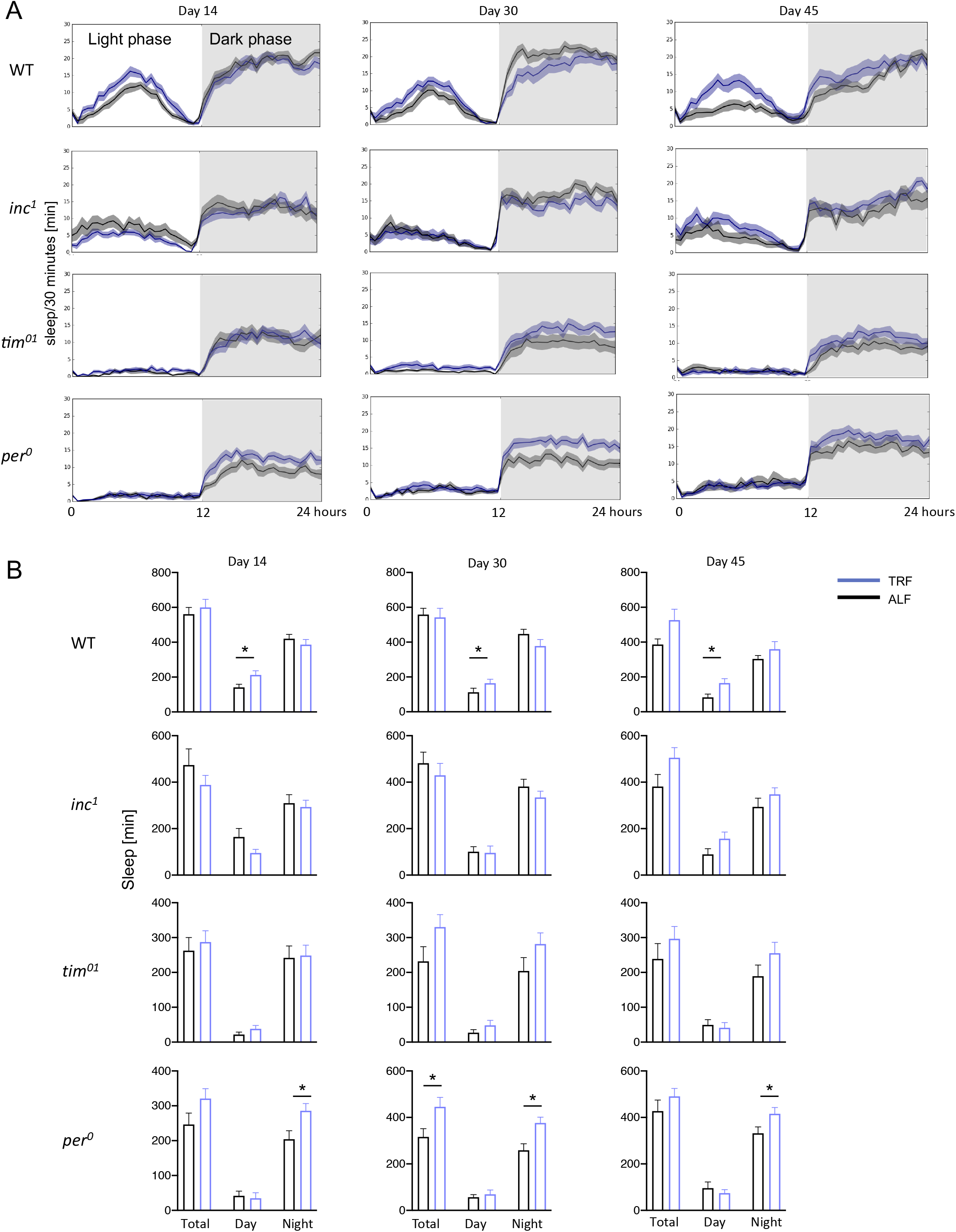
TRF and sleep. Female wild-type, *inc*^*1*^, *per*^*0*^ and *tim*^*01*^ flies were treated with TRF or ALF. Sleep was assessed after 14, 30 or 45 days. (A) For all sleep plots, average 24 h sleep over 4 days binned as sleep in 30 min is shown. (B) Quantification of (A). Shown are bar charts of mean total daily sleep, daytime sleep and nighttime sleep as well as SEM. 16 flies were tested per condition. t-tests were used to assess significance. *=P≤0.05

**Supp. Fig 3.**
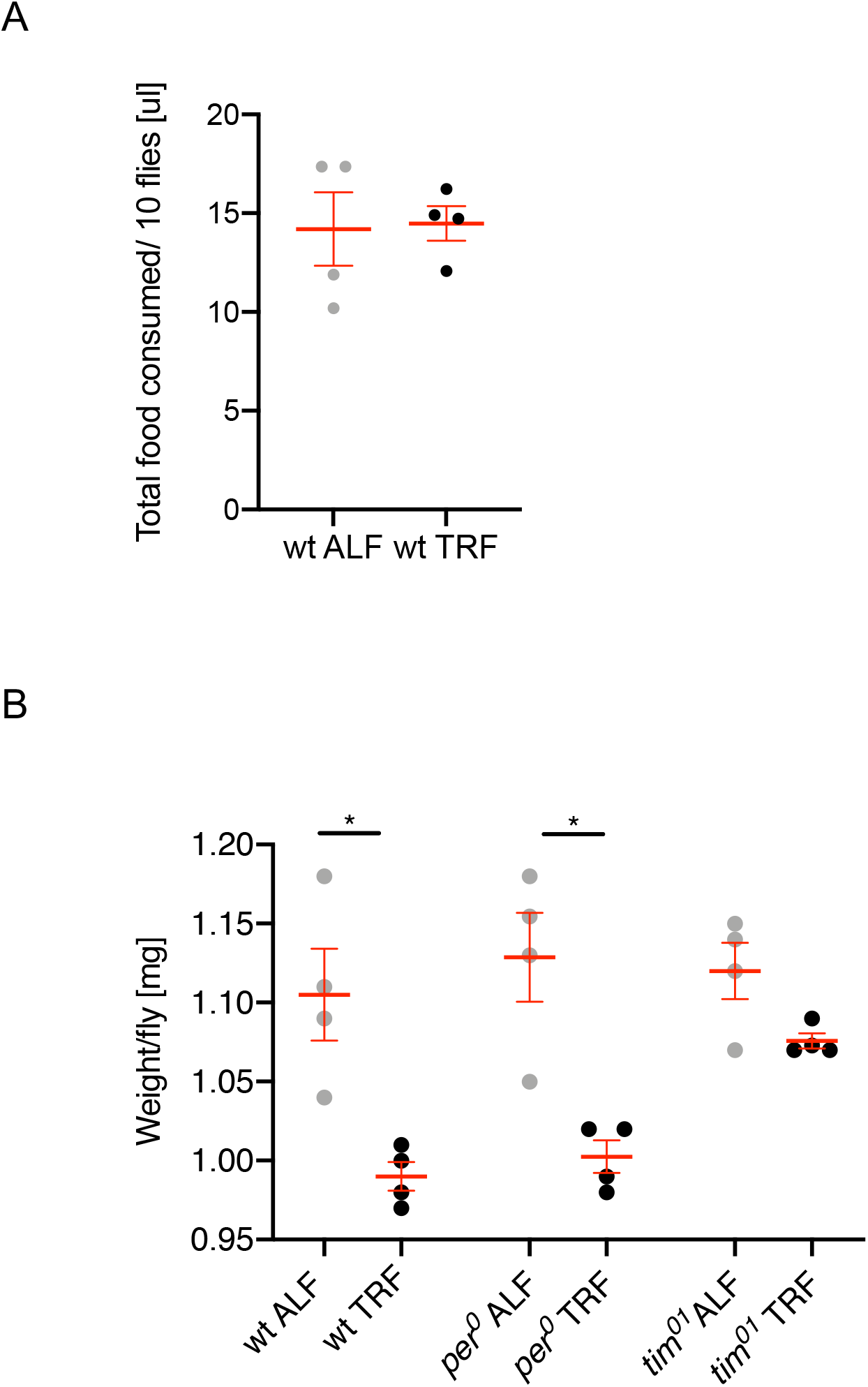
Impact of TRF on feeding. (A) Female wild-type flies underwent 32 days of TRF or ALF in 12:12 LD. Food intake was measured in groups of 10 flies for 12 hours (TRF group) or 24 hours (ALF group) using the capillary feeder assay (CAFE). Each point represents food consumption of 10 flies. (B) Female wt, *per*^*0*^ and *tim*^*01*^ flies underwent 45 days of TRF or ALF in 12:12 LD and were weighed in groups of 10 at ZT 10. Each point represents the weight of 10 flies divided by 10. Mean and SEM are indicated in red. Mann-Whitney testing assessed statistical significance. *=P≤0.05

**Supp. Fig 4.**
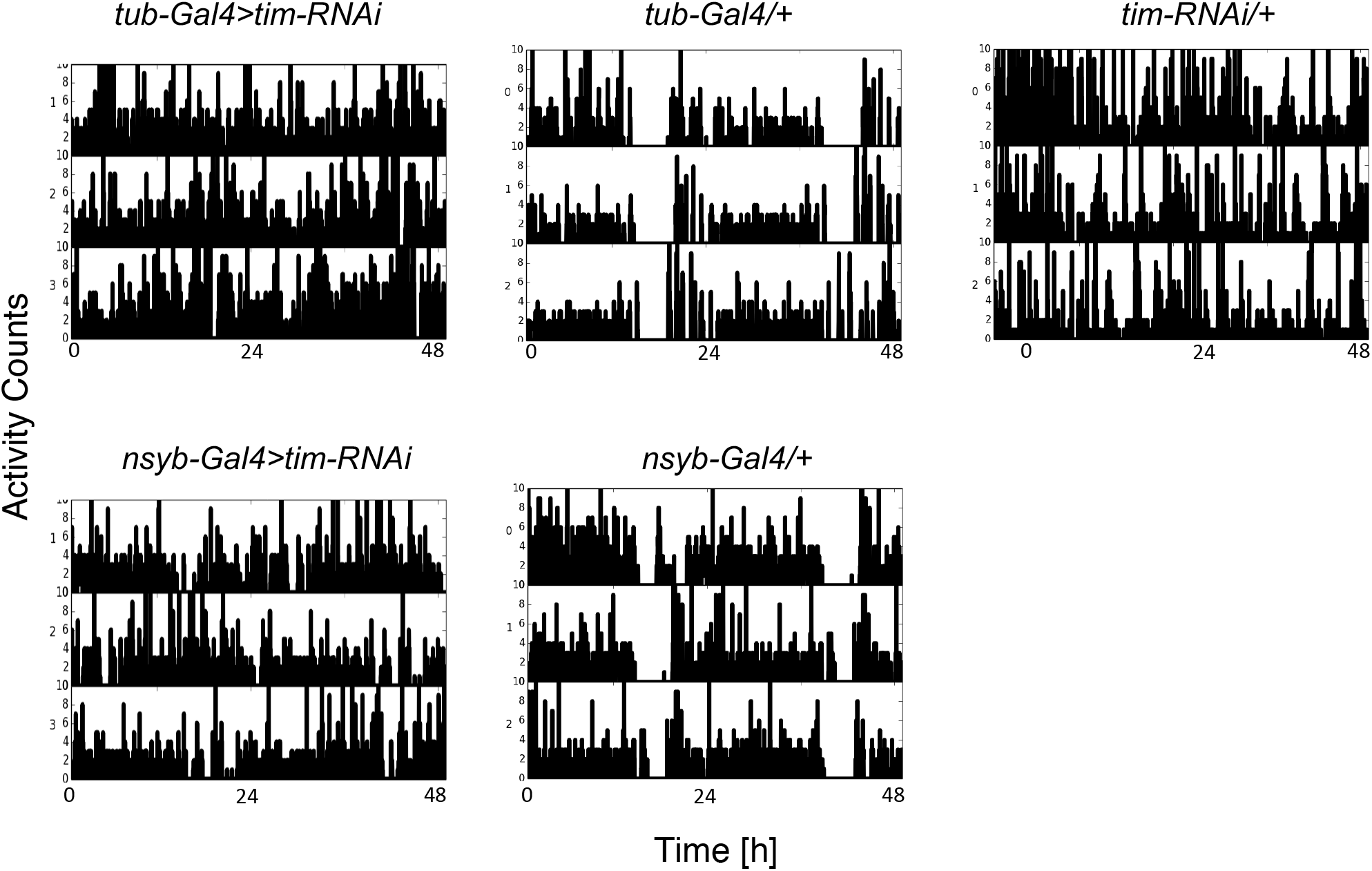
Free-running activity of flies with *tim* knockdown in different tissues. 2-3 day old flies of the indicated genotypes were entrained in LD cycles for 3 days and moved to DD to assess rhythmicity. Ubiquitous *tim-RNAi* using *tub-Gal4* as well as neuronal knockdown using *nsyb-Gal4* causes arrhythmicity, while heterozygous controls remain rhythmic. Shown are representative examples of double-plotted activity counts of individual flies. In contrast to male flies, in our hands mated female *Drosophila* of all genotypes show a higher proportion of arrhythmic flies, which might be due to reduced sleep and other previously described sexual dimorphisms in *Drosophila* locomotor behavior (See discussion).

